# DNA sliding and loop formation by *E. coli* SMC complex: MukBEF

**DOI:** 10.1101/2021.07.17.452765

**Authors:** Man Zhou

## Abstract

SMC (structural maintenance of chromosomes) complexes share conserved architectures and function in chromosome maintenance via an unknown mechanism. Here we have used single-molecule techniques to study MukBEF, the SMC complex in *Escherichia coli*. Real-time movies show MukB alone can compact DNA and ATP inhibits DNA compaction by MukB. We observed that DNA unidirectionally slides through MukB, potentially by a ratchet mechanism, and the sliding speed depends on the elastic energy stored in the DNA. MukE, MukF and ATP binding stabilize MukB and DNA interaction, and ATP hydrolysis regulates the loading/unloading of MukBEF from DNA. Our data suggests a new model for how MukBEF organizes the bacterial chromosome *in vivo*; and this model will be relevant for other SMC proteins.

## Introduction

The SMC (structural maintenance of chromosome) complexes share conserved architectures and function in chromosome maintenance throughout all kingdoms of life, including condensin, cohesin, and SMC5/6 in eukaryotes and MukBEF in *E. coli* (*1*) (*2*) (*3*). SMC dimers adopt a large ring structure containing a ~50-nm antiparallel coiled-coil “arm” with a hinge dimerization domain and an ABC-type ATPase head domain (*4*). The distinctive and conserved molecular architecture suggests a common principle of SMC complex function. One mechanism explaining the formation of DNA loops is provided by the loop extrusion model, which proposes SMC proteins to act as loop-generating motors during iterative cycles of ATP hydrolysis. Single molecule assays have demonstrated that condensin and cohesin translocate along DNA in an ATP-dependent manner (*5*) (*6*); in contrast, other studies have reported that cohesin slides along DNA in an ATP-independent manner (*7*) (*8*). Meanwhile, there are reports of real-time visualization of symmetric/asymmetric DNA loop extrusion by condensin and symmetric loop extrusion by cohesin (*9*) (*10*) (*11*) (*12*).

However, all proposed models ignore the physical nature of DNA as a highly dynamic polymer with properties that may profoundly affect the function of SMC proteins. Overlap of two DNA polymers in cylindrical confinement significantly reduces their conformational entropy, and theoretical simulations suggest that entropic forces can drive chromosome segregation under the right physical condition. (*13–17*) Moreover, experimental studies have already well characterized the *E. coli* nucleoid dynamics in living cells (*18*) (*19*). Another simulation-based model suggested that chromatin loops can be efficiently formed by non-equilibrium dynamics of DNA on the diffusive sliding of molecular slip links (*20*).

Here, we exploit two complementary single-molecule assays with total internal reflection fluorescence (TIRF) microscopy to probe the function of the *E. coli* SMC complex, MukBEF, by either immobilizing DNA or MukB on the surface (Fig.1B).

**Fig.1.**
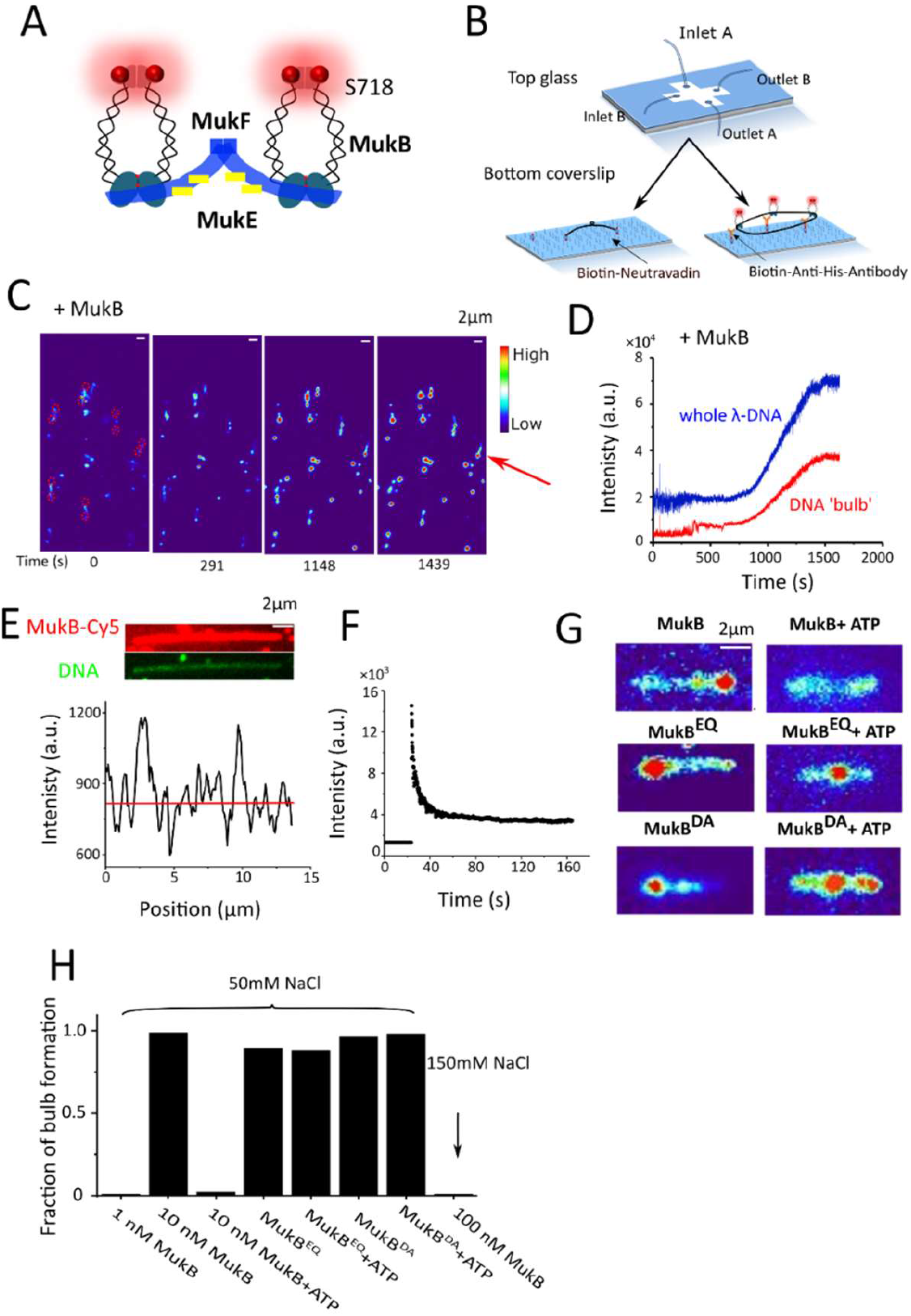
DNA compaction by MukB in the absence of ATP. (A) Cartoon representation of MukBEF subunits and domains in different colors. MukF (blue) is kleisin protein which binds and bridges the ATPase head domains of the MukB dimers, and MukE (yellow) is KITE protein in *E. coli*. S718 at hinge domain was mutated to unnatural amino acid (p-azido-l-phenylalanine, AZF) and labelled with Cy5 (marked as red balls) via click chemistry. (B) Sketch of single molecule study. Top slide is designed for injection along two perpendicular directions. The bottom slide is to immobilize either MukB or DNA on the surface. In assay 1, 48.5 kbp *λ*DNA was doubly tethered to the PEG modified surface. In assay 2, MukB was immobilized by using a His_6_-tagged variant of MukB and biotinylated anti-His_6_-antibody on the PEG surface. (C) Time course showing DNA compaction by 10 nM MukB at 50 mM NaCl. DNA ‘bulbs’ are visible as bright dots on the doubly tethered *λ*DNA. Two tethered ends are marked by red dashed circles. The red arrow shows one example of a doubly tethered *λ*DNA (D) The fluorescence intensity growing over time for whole *λ*DNA (blue curve) and for DNA ‘bulb’ (red curve) as indicated by red arrow in Fig. 1C. The distance between two tethered ends is ~4 μm. We noticed that it was hard to quantify the compaction rate, as in TIRF microscope, the laser intensity decays exponentially with increasing distance from the interface; therefore, the intensity of entire *λ*DNA is low at the beginning as most *λ*DNA fragments are floating out of TIRF field. As time goes, more DNA is compacted coming close to the surface, and the increased intensity of ‘DNA bulb’ (Fig. 1D) comes from two fractions: one from compacted DNA, and the other from the increased laser intensity close to the surface. (E) Distribution of Cy5 labelled MukB on over-stretched DNA (DNA curtain; video S5). The top panel shows two spectral channels: the red one is a snapshot of Cy5 labelled MukB, and the green one is a snapshot of Sytox orange stained DNA. The bottom panel is the fluorescence distribution of Cy5 along the DNA. (F) Bleaching curve of Cy5 in Fig. 1E over time, which indicates that many MukB molecules binding to DNA. (G) Snapshot examples of *λ*DNA under different conditions. DNA compaction and DNA ‘bulb’ formation is induced by MukB only without ATP. However, ‘bulbs’ can be seen after addition of MukB^EQ^ and MukB^DA^, both in the absence and presence of ATP. Scale bar 2 μm. (H) Fraction of DNA bulbs that formed on doubly tethered *λ*DNA following incubation with the indicated components at different salt conditions. 50-150 DNAs were analysed.

## Results

### DNA compaction by MukB at low salt in the absence of ATP

Condensin and cohesin can compact DNA in the presence of ATP (*9–11*) (*21*). However, in assay 1 (Fig. 1B), no DNA compaction can be observed on the doubly tethered *λ*DNA while incubation with MukBEF and ATP-Mg^2+^ (Video S1). Fluorescently labelled MukB (MukB-Cy5) shows there is no MukBEF stable association with DNA during the observation time (Video S2). Importantly, fluorescent labelling does not affect MukBEF ATP hydrolysis activity (Fig.S2).

To examine the function of MukBEF carefully, we tested the MukBEF components sequentially. First, we incubated 10 nM MukB alone with DNA in the absence of flow, all slack *λ*DNA molecules became tight **(compaction)** (Fig. 1C, video S3), and DNA ‘bulbs’ **(locally compacted DNA)** can be observed frequently (98.7%, 149/151 of DNAs, and Fig. 1H). DNA compaction is gradual over time (Fig. 1C and 1D) and the compaction rate is MukB concentration dependent, with higher compaction rate at higher concentration of MukB. For example, 10 nM MukB takes ~1000 s to fully compact DNA, while 100 nM MukB takes ~150 s (video S3 & video S4). We noticed that there is a ‘threshold’ protein concentration and salt sensitivity required for DNA compaction. For example, at 50 mM NaCl, 1 nM MukB alone cannot compact DNA, and ‘bulbs’ are rarely observed even after 4 hours (1.7 %, 1/60 DNAs), while for concentrations larger than 10 nM, MukB can easily compact DNA and ‘bulbs’ are frequently observed (98.7%, 149/151 of DNAs). At 150 mM NaCl, even 100 nM MukB cannot compact DNA and ‘bulbs’ are rarely observed (0 %, 0/58 DNAs, Fig. 1H). This compaction is not stable and can be easily destroyed by resuming high flow rates (> 0.5 pN), and that DNA ‘bulbs’ cannot be expanded into loops at ~1 pN of applied force.

To test whether MukB has a sequence preference for prominent loading sites to form ‘bulbs’, we over-stretched the DNA (similar as DNA curtain) and added MukB-Cy5; a fairly homogenous distribution of MukB along the DNA can be observed (Fig. 1E & video S5), consistent with MukB loading randomly onto DNA. The bleaching curve of MukB-Cy5 indicates multiple MukB molecules binding to DNA (Fig. 1F).

### ATP inhibits MukB-mediated DNA compaction in the absence of MukEF

We further tested the function of ATP during DNA compaction by MukB. Strikingly, DNA cannot be compacted by MukB incubation with ATP-Mg^2+^ and no ‘bulbs’ can be seen on DNA (the fraction of DNA bulbs: 1.0 %, (1/ 98 of DNAs) Fig. 1G & Fig. 1H). Meanwhile, the dwell time of MukB on DNA in the presence of ATP-Mg^2+^is very short and stable association of MukB with DNA cannot be observed (Video S6).

Constructs with EQ mutation in the SMC ATPase domain, where the two catalytic domains remain in an ATP engaged condition, are deficient in ATP hydrolysis and exhibit abnormally stable binding to DNA (*22*) (*23*) (*24*). *In vivo* studies suggest that an EQ mutation holds together newly replicated chromosomes and inhibits chromosomes segregation (*25*), or inhibits the relocation of SMC proteins (MukBEF and cohesin) (*26*) (*27*). We further studied the EQ MukB mutant (MukB^EQ^), which can bind but cannot hydrolyse ATP. Strikingly, MukB^EQ^ can fully compact DNA both in the presence and absence of ATP-Mg^2+^ at a MukB concentration above 10 nM (fraction of DNA bulbs: 90.2 %, (46/51 of DNAs) and 88.0%, (103/117 of DNAs) Fig. 1G, Fig. 1H and video S7 & 8). Another DA mutation (MukB^DA^) which cannot bind ATP, can also fully compact DNA both in the presence and absence of ATP-Mg^2+^ at a higher MukB concentration (100 nM) (fraction of DNA bulbs: 96.3 %, (52/54 of DNAs) and 98.1%, (51/52 of DNAs) Fig. 1G, Fig. 1H and video S9 & 10). The EQ and DA mutation data suggest that ATP does not contribute directly to DNA compaction.

### MukEF and ATP-Mg^2+^conditionally stabilize pre-incubated MukB and DNA complex

MukEF depleted strains show the same temperature-sensitive growth phenotype as the MukB-null strain, and MukEF is essential for stable association of MukB with chromosome *in vivo* (*28*). Thus, we further investigated the function of MukEF and ATP-Mg^2+^, for which we first incubated MukB with DNA until DNA is fully compacted, then flowed in MukEF with ATP-Mg^2+^. We observed that DNA just becomes slack, and stable ‘bulbs’ can be observed (Fig. 2A) under high flow. Occasionally, these ‘bulbs’ can be expanded into ‘loops’, however, these loops do not grow in size over time. (Fig.2B & video S12). Most MukB was just washed off, with some clusters remaining on DNA, and quantification shows that the numbers of MukB varies. 84.1% (Fig. 2D) measured MukB shows the number between 2 to 4, which indicates that the major population of MukBEF clusters are dimers of homodimeric MukB. These ‘bulbs’ are very stable and remain associated with DNA even under ~2.5 pN of applied force.

**Fig. 2.**
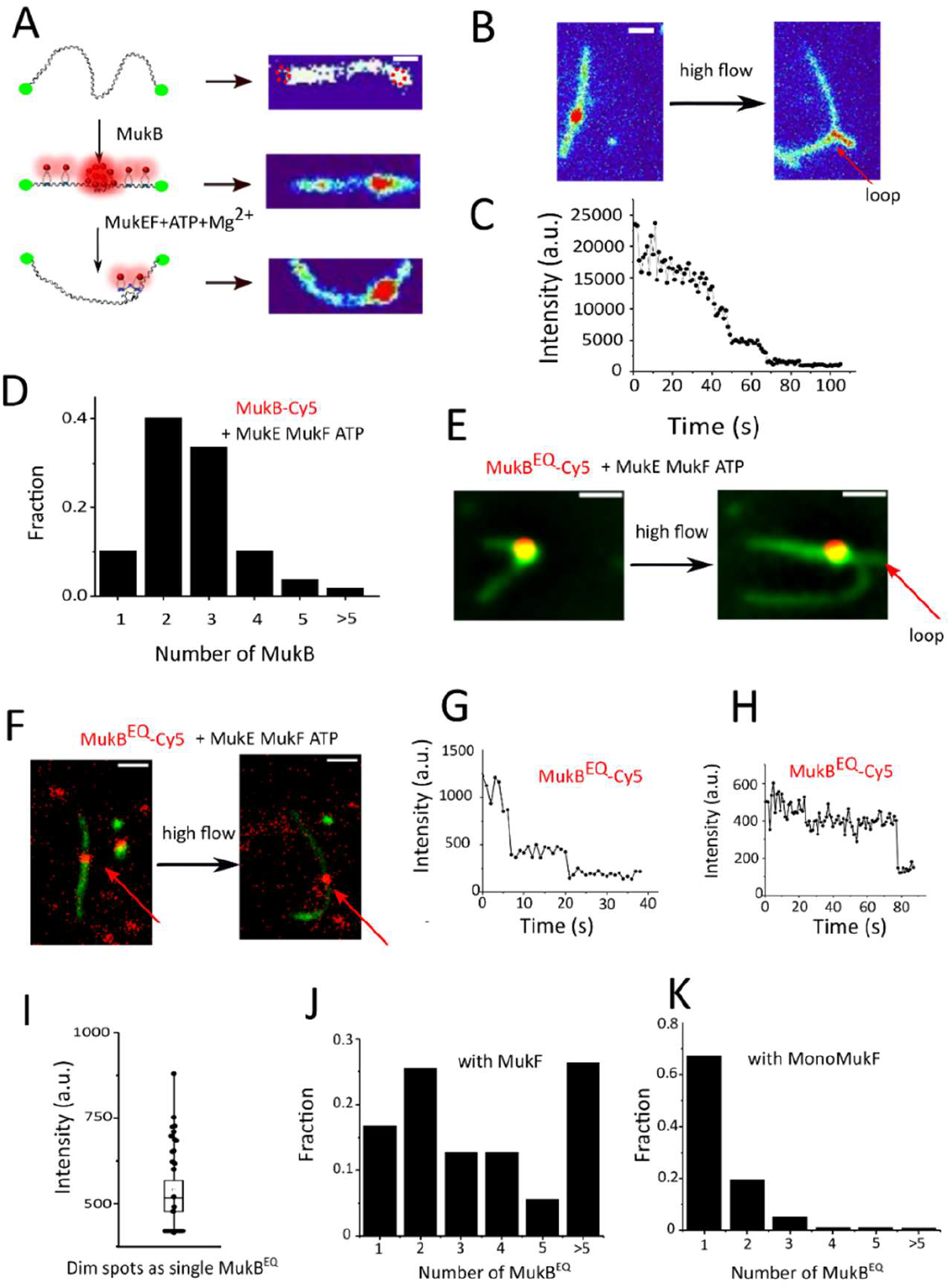
DNA compaction and loop formation under different conditions. (A) DNA ‘bulbs’ on the u-shaped double-tethered *λ*DNA by MukBEF clusters with ATP-Mg^2+^. First, *λ*DNA was incubated with MukB until DNA is fully compacted, and afterwards, MukEF and ATP-Mg^2+^ were flowed into the chamber from a perpendicular direction. Scale bar 2 μm. The right panels show examplary snapshots of different DNA conformations. (B) One example of a ‘bulb’ formed in the presence of MukBEF clusters with ATP-Mg^2+^. This ‘bulb’ can be expanded into loops at ~1 pN of applied force. The loop does not grow over time (Video S12). Scale bar 2 μm. (C) Representative time traces of fluorescence intensities of MukBEF clusters with ATP-Mg^2+^on tethered DNA. (D) The chart shows numbers of MukB clusters with ATP-Mg^2+^on DNA ‘bulbs’ based on measuring the fluorescence intensity of the ‘bulbs’ and comparing it to that of a single MukB bound to glass (bleaching steps are too noisy for reliable quantification). Quantification shows 40.2% of 2-MukB, 33.6% of 3-MukB, and 10.3% of 4-MukB molecules; 107 DNAs were analysed. (E) Stable ‘bulbs’ and ‘loops’ can be easily observed when MukB^EQ^, MukEF and ATP-Mg^2+^are present. One example of a ‘bulb’ that can be expanded into a loop (red arrow) at ~1 pN of applied force (Video S14). Scale bar 2 μm. (F) Stable association of MukB^EQ^ can be easily observed on DNA at a decreased concentration of MukB^EQ^(1 nM), MukE2F (2 nM) in the presence of 1 mM ATP-Mg^2+^. However, no ‘bulbs’ or ‘loops’ can be observed under flow. Labelled MukB^EQ^ (marked by red arrow) moves together with DNA, confirming that it was indeed stably bound to DNA. Scale bar 2 μm. (G) Representative two stepwise bleaching events of MukB^EQ^ in the presence of MukEF and ATP-Mg^2+^. (H) Representative single step bleaching of MukB^EQ^ in the presence of MukEF and ATP-Mg^2+^. (I) Intensity distribution of single MukB^EQ^ EF clusters on the surface. (J) The chart shows numbers of MukB^EQ^ on DNA ‘bulbs’ in the presence of MukEF and ATP-Mg^2+^, based on measuring the fluorescence intensity of the ‘bulbs’ and comparing it to that of single MukB^EQ^ bound to glass. 125 clusters were analysed. (K) The chart shows numbers of MukB^EQ^ on DNA ‘bulbs’ in the presence of MukE, MonoMukF, and ATP-Mg^2+^. 164 clusters were analysed.

### Stable loop formation by MukB^EQ^EF and ATP-Mg^2+^

Because MukB^EQ^EF can form clusters *in vivo* (*26*), we wanted to test the behaviour of MukB^EQ^EF *in vitro*. Here, we observed that MukB^EQ^EF forms clusters on the DNA with MukB^EQ^ EF and ATP-Mg^2+^, and these MukB^EQ^EF complexes are very stably associated on DNA with 150 mM NaCl buffer and at ~1 pN of applied force (Fig. 2E, Fig. S5A & video S13). DNA ‘bulbs’ can also be observed, and frequently these ‘bulbs’ can be expanded into ‘loops’ at ~1 pN of applied force, however, the loops don’t grow in size over time even when applying ~1 pN force (Fig. 2E & video S14). Quantification shows that the number of MukB^EQ^EF varies (Fig. 2J), and higher-number clusters appear more frequently compared to the wt-MukBEF with ATP-Mg^2+^ (Fig. 2D and Fig. 2J). When we decreased MukB^EQ^ to 1 nM, stable association of MukB^EQ^ with DNA was also observed, however, without DNA ‘bulbs’ (Fig. 2F). We speculate that these DNA ‘bulbs’ and ‘loops’ are formed by dimerization of MukF, because they are very stable against high flow rates, in contrast to DNA ‘bulbs’ formed by MukB alone and which are not stable under high flow rates. To test this, we flowed in MukB^EQ^ with truncated MukF (Monomeric MukF, MonoMukF) which is monomeric in solution, being unable to form stable heads-engaged dimers (*29*). Very stable protein complexes can be observed on DNA with MonoMukF with 150 mM NaCl buffer and at ~1 pN of applied force (Fig. S5B and video S15); however, DNA ‘bulbs’ or ‘loops’ were rarely observed. Quantification shows that 88.4 % of cluster population contains less than 2 MukB^EQ^, indicates MukB^EQ^ is a single dimer with MonoMukF. (Fig. S4 & Fig. 2K).

### DNA can be stretched on immobilized MukB and slides through MukB

To further investigate the dynamics of DNA under action of of MukBEF, in assay 2, MukB was immobilized on the surface, and the interaction of DNA with MukB(EF) was analysed (Fig. 1B). First, we pre-incubated circular DNA (44kb plasmid) with surface-immobilised MukB-His, as well as MukE-Flag, MukF-Flag and ATP-Mg^2+^ in solution; in this case, essentially no DNA spots were observed on the surface (Fig. 3A and video S16). However, upon incubation of MukB-His with plasmid DNA, many DNA spots can be observed on the surface (Fig. 3A & video S17). For the control, we barely observed any DNA spots on the biotinylated anti-His_6_-antibody functionalized PEG surface in the absence of MukB-His, which indicates that the imaged DNA molecules are specifically captured by MukB. Under flow, we observed stretched plasmids on the surface and breaking of DNA, with laser excitation commonly triggering DNA sliding off the surface (Fig. 3B & video S18).

**Fig. 3.**
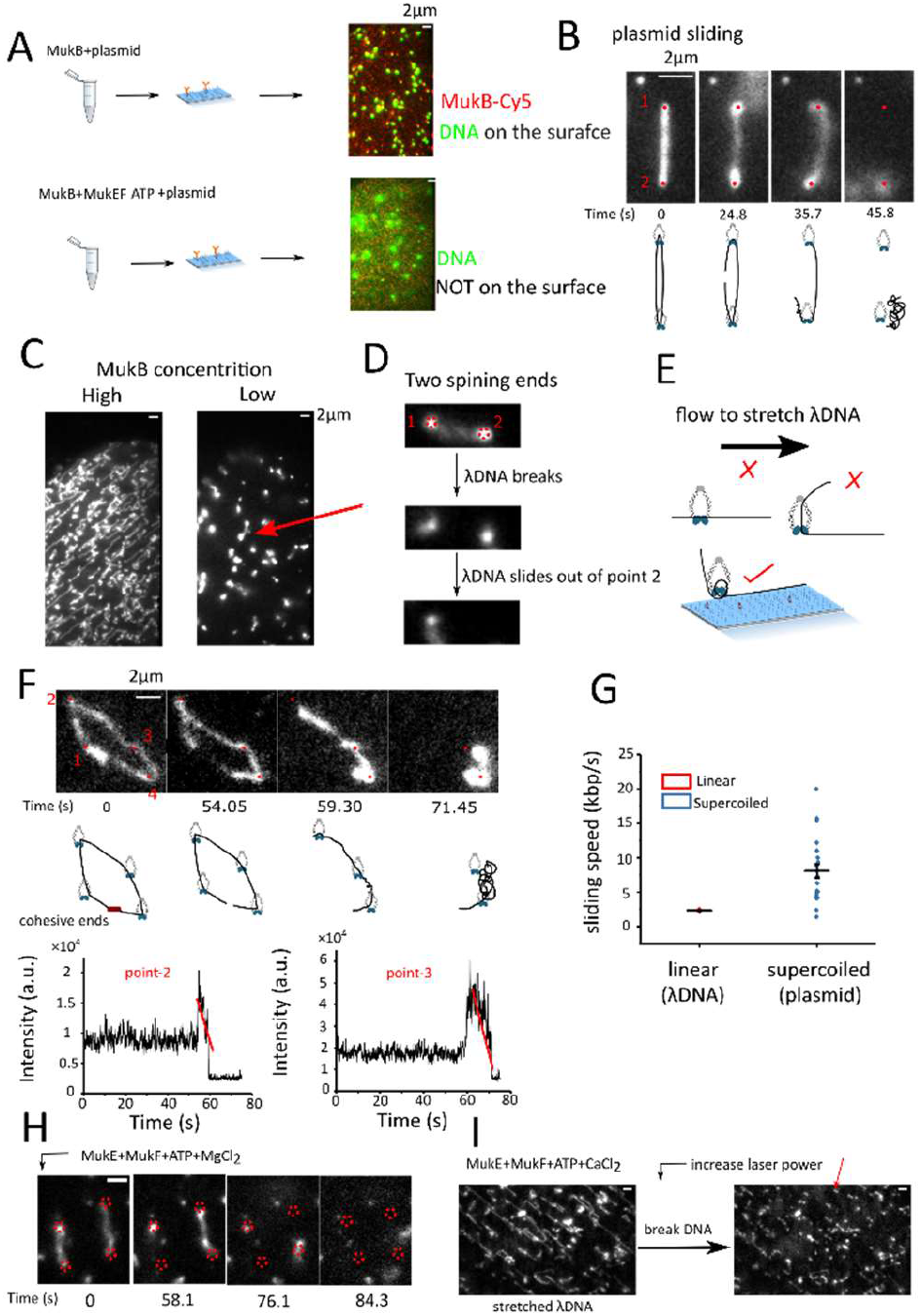
Immobilization of His_6_-tagged MukB on the surface. (A) Upper panel: example of circular DNA (44 kb plasmid, green, Sytox orange staining) captured on the surface and incubated with His_6_-tagged MukB. Lower panel: example of circular DNA diffusion in solution incubated with 10 nM MukB with 2mM ATP+ 2 mMCaCl_2_, +10 nM MukE + 20 nM MukF (B) Sequential images of 44 kb plasmid sliding through MukB. Two red dots indicate the tether points. At ~24.8 s, the plasmid DNA breaks. At ~35.7 s and ~45.8 s, the plasmid DNA slides through the tether points one after the other. (Video S18) (C) Linear DNA (*λ*DNA) can be stretched on surface with immobilized MukB. Left panel: 150 pM *λ*DNA with 20 nM MukB, right panel: 150 pM *λ*DNA with 5 nM MukB. Incubation buffer: 10 mM NaCl 20 mM Tris-HCl pH=7.0. (D) One example with two DNA ends spinning round the tether points (Video S19). Red dashed circles indicate two captured points on the surface with immobilized MukB. DNA breaks and slides out of point-2. (E) *λ*DNA topologically entrapped in the MukB ring can be stretched along the surface. (F) *λ*DNA slides through immobilized MukB and is sequentially released. Sequential images of *λ*DNA sliding across MukB. Series of snapshots showing Sytox orange-stained *λ*DNA sliding through tethered MukBs one by one (Video S20). Four red dots indicate the tether points. At ~54 s, the cohesive end breaks. At ~59.3 s and ~71.45 s, the *λ*DNA slides through the second and the third tether points, respectively. The DNA slides in a clockwise direction. Notably, the *λ*DNA does not slide through the fourth tether point in anti-clockwise direction. Schematic diagrams under each snapshot are for visual guidance. Two plots monitor the DNA intensity around the tether points (point 2 and point 3) over time. The linear decay (red lines) indicates that the DNA release mechanism is by sliding, instead of multiple binding/unbinding. (G) Quantification of sliding speed of linear DNA (*λ*DNA) and supercoiled DNA (44 kb plasmid). (H) Time course of captured *λ*DNA on surface with immobilized MukB under high salt wash (150 mM) with MukE, MukF and ATP-Mg^2+^. The capture points are marked by red dashed circles. One capture point was released from surface at 58.1 s, and two other points were released at 76.1 s, and the last capture point was released at 84.5 s. (video S21) (I) Captured *λ*DNA on surface with immobilized MukB under high salt wash (150 mM) with MukE, MukF and ATP-CaCl_2_ (video S22). Left panel: *λ*DNA is totally stretched on surface with the immobilized MukB. Right panel: laser excitation induces *λ*DNA break, and DNA debris is still at the tether points on the surface. One examplary tether point is marked by red arrow.

To test what happens to linear DNA, we first incubated MukB with *λ*DNA at 50 mM NaCl, then flowed the mixture into the cell functionalized with biotinylated anti-His_6_-antibody on the surface. Strikingly, *λ*DNA can be easily stretched on the surface with many tether points and two DNA ends spinning round the tether points (Fig. 3C). At low concentration of MukB, *λ*DNA was stretched only with two tether points, and laser-induced breaking of DNA commonly triggers DNA sliding off the tether points (Fig. 3C, Fig. 3D & video S19). Stretching DNA normally requires modification of DNA extremities to anchor DNA to a functionalized substrate, e.g., via sticky ends or biotin functionalization, and the bond strength between biotin and streptavidin is far beyond the weak electrostatic interaction between MukB and DNA (Fig. S3). Why could naked *λ*DNA be stretched on the surface? One possibility is that the DNA is topologically entrapped in the MukB ring (Fig. 3E), which inhibits DNA detachment from the surface even under strong flow.

For open *λ*DNA, it is very difficult to estimate the DNA length and thus to calculate the sliding speed, as many DNA fragments are floating out of the TIRF region. We managed to get two circular *λ*DNA. After DNA breakage, *λ*DNA slides on immobilized MukB and sequentially detaches (Fig. 3F & video S20). We further calculated the sliding speed for *λ*DNA, which was ~2kbp/s (Fig. 3G). For plasmids, the sliding speed was ~5kbp/s, faster than for linear *λ*DNA (Fig. 3G).

### MukE, MukF and ATP binding stabilizes MukB and DNA complex on the surface

To further investigate the function of MukEF and ATP-Mg^2+^, we then flowed in MukE-Flag, MukF-Flag and ATP-Mg^2+^ in 150 mM NaCl buffer to investigate the behaviour of *λ*DNA on tethered MukB; after these additions, most *λ*DNA molecules were just washed off (Fig. 3H & video S21). When we flowed in MukE-Flag, MukF-Flag and ATP-Ca^2+^, which is supposed to inhibit ATP hydrolysis of the nucleotide-binding domains (*30*), strikingly, most *λ*DNA remained on the surface. We then significantly increased the flow rate up to 100ul/min (~ 2.5 pN) with MukE-Flag, MukF-Flag and ATP-Ca^2+^ in 150 mM NaCl, and all *λ*DNA were over-stretched on the surface to linear conformations with a few tether points (Fig. 3I & video S22). We then increased the laser power to break DNA into pieces (Fig. 3I & video S22). Strikingly, *λ*DNA does not slide off the surface, and the tether points remained on the surface. This suggests that without ATP hydrolysis, MukE, MukF and ATP stabilize the interaction between MukB and DNA, and this stabilized interaction can even resist high force (~ 2.5 pN).

## Discussion and conclusions

Our results show that MukB alone can compact DNA under low salt conditions when MukB concentration is above a certain ‘threshold’ concentration. This is in agreement with previous studies that showed MukB alone can compact DNA (*31*) (*32*). This ‘threshold’ concentration is easily explained by conventional DNA-protein interaction kinetics as shown in Fig.4 A. It should be noted that this *in vitro* observation cannot be tested *in vivo*, as ATP is always present in a cell.

As MukB dimers alone have negligible ATPase activity (*33–35*), the catalytic cycle of ATP hydrolysis may influence the dwell time of MukB on DNA by regulating the loading/unloading of MukB on and from DNA, when MukE and MukF are absent, and this could be the underlying reason why ATP inhibits MukB-mediated DNA compaction. MukB, ATP-Mg^2+^, without MukEF shows no DNA compaction *in vitro* which is in agreement with *in vivo* observation that MukEF is essential for stable association of MukB with DNA, and no MukB stable clusters were detected in cells deficient in MukE or MukF (*36*), (*28*).

In this study, we showed that MukB^EQ^ and MukB^DA^ does also compact DNA under low salt conditions. Our results show that MukB^EQ^ with MukE, MukF and ATP-Mg^2+^ can form stable complexes on DNA. These complexes are extremely stable even at 150 mM NaCl and high flow rates. DNA ‘bulbs’ and ‘loops’ can be observed, and these loops are more likely formed by dimerization of MukF, as loops were not observed with MonoMukF. Loading and unloading of MukB^EQ^EF on/from DNA can easily be elucidated by the binding kinetics of MukB^EQ^EF and ATP (Fig. 4D).

**Figure 4:**
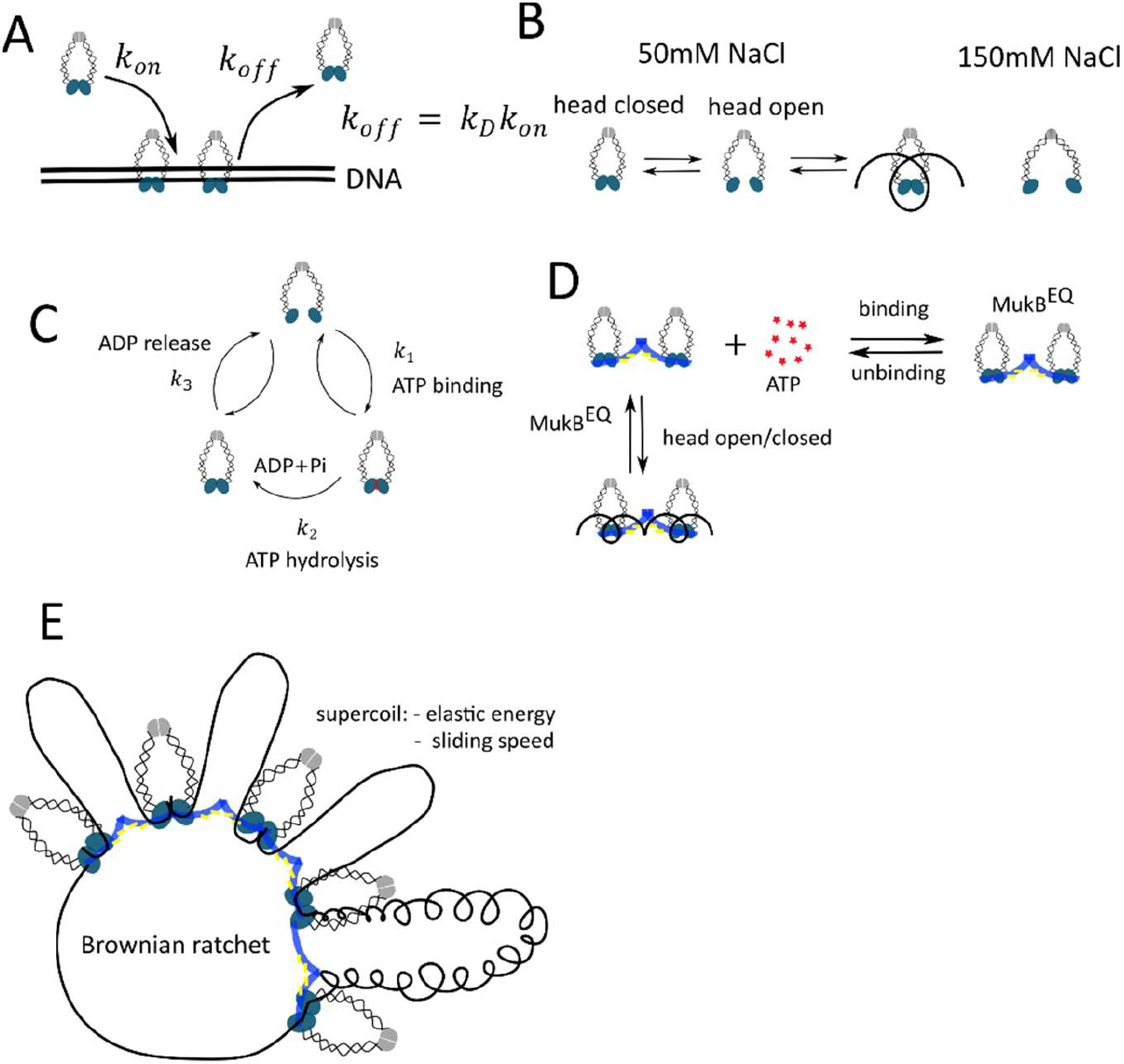
Kinetics of MukBEF loading/unloading on/from DNA and Brownian ratchet model. (A) Binding kinetics of MukB and DNA. *k_off_* and *k_on_* represent dissociation (off-rate) and association (on-rate) rates of MukB from to/ DNA, respectively. *K_D_* represents binding affinity between MukB and DNA. The balance of on and off rates determines the binding occupation of MukB on the DNA. As off-rate *k_off_* is usually assumed to be independent of protein concentration, the MukB concentration has to be above a ‘threshold’ concentration to gradually decorate DNA for inducing the observed DNA compaction. (B) Equilibration between closed and open head states at 50mM NaCl. We assume that low salt concentration (50 mM NaCl) artificially increases the stability of the head domain. At this salt concentration, the MukB head is transiently opening the closed MukB ring due to thermal fluctuations, which allows DNA to be topologically entrapped in the MukB ring and potentially wrapped around the head domains. In contrast, under high salt concentration (150 mM NaCl), no stable interaction between MukB and DNA can be observed. (C) Kinetics of ATP hydrolysis cycle. The mean turn-over time of ATP hydrolysis measured in a test tube consists of three contributions: ATP binding rate (*k*_1_), single ATP turn-over rate (*k*_2_), and ADP releasing rate (*k*_3_). We assumed that the ‘real’ ATP hydrolysis rate (*k*_2_) can be regarded to be constant at physiological concentrations of ATP/ADP. ATP binding (*k*_1_ requires that two head domains are very close to each other, which is the rate-limiting step of the ATPase cycle. When only MukB is present, the ATPase head domain does freely diffuse and it takes comparably long time to bind ATP (slow *k*_1_), in which case one observes negligible ATPase activity. MukF promotes contact between MukB heads by restricting their free diffusion, which increase the ATP binding rate (*k*_1_, and therefore leads to increased ATPase activity. MukEF inhibits MukB head rotation/diffusion for ADP release, therefore, a decrease of *k*_3_ and a slightly decreased ATPase activity as compared to MukF is observed. Overall, MukF and MukE regulate ATP binding /ADP releasing, leading to the observed ATP hydrolysis rates in the test tube. (D) Binding kinetics of MukB^EQ^EF, ATP and DNA. MukB^EQ^EF without ATP is capable of opening the MukB ring at a low possibility due to thermal fluctuations, which allows DNA entrapment. As ATP hydrolysis regulates MukB head opening/closing. Without ATP hydrolysis, MukB^EQ^EF cannot be unloaded from DNA, therefore, MukB^EQ^EF is very stably associated on DNA, even under 150 mM NaCl and ~1 pN force. (E) Brownian ratchet model for DNA sliding through MukBEF. The big loops are formed by dimerization of MukF. DNA sliding through MukB regulates loop size. Supercoiling of DNA, which is related to the elastic energy of DNA, may regulate the sliding speed.

There exists one core argument that MukBEF together with ATP-Mg^2+^ all are required for normal chromosome organization and segregation *in vivo*; while our *in vitro* single-molecule data shows that MukBEF together with ATP-Mg^2+^ seems to be incapable of remodelling DNA; only rare MukBEF clusters on DNA ‘bulbs’ were observed occasionally. Actually, this is in agreement with *in vivo* observations that even high concentrations of ~100 nM of (Muk4B:4E:2F)-complexes at high density of DNA in *E.coli* (the density of DNA in *E.coli* is 10^5^ times of DNA used in the single-molecule assay) result only in 48% loading on DNA with a dwell time of about ~60 s. The unloading rate (*k_off_MukBEF_*) is ~0.015 s^-1^ (dwell time 65 s), and the loading rate *in vivo (k_on_MukBEF_*) is only ~3.9*10^-6^ s^-1^ which would result in 48% occupancy of MukBEF on chromosome (*26*) (*37*). In the single-molecule assay, it requires 1/3.9*10^-6^ = 2.56*10^5^ s (~ 3 days) to capture a loading event in pre-incubated MukBEF solution, and that is also the reason why in an *in vitro* biochemistry assay which shows a ‘fake’ inhibition of MukEF to loading of MukB on DNA. The extremely low loading rate does also reveal that most collisions of MukBEF with DNA do not lead to loading, and there is a small chance that ATP binding or hydrolysis can trigger distinct conformational changes within the two head regions of MukBEF that could act synergistically to open the MukB ring for DNA loading/unloading. ATP hydrolysis is required to regulate MukB head opening/closing, and therefore regulates MukBEF loading/unloading on/from DNA to balance the numbers of MukBEF on DNA, as too many loops generated by MukBEF lead to over-compacted chromatin which may block chromosome replication, gene expression and regulation, whereas too few generated loops lead over-spread chromatin.

Pre-incubated MukBEF complexes rarely undergo stable association with DNA, while sequential incubation of MukB with DNA, and then MukEF and ATP-Mg^2+^, can stabilize MukB/DNA interaction. Together with *in vivo* observations that MukEF is essential for stable association of MukB, these superficially conflicting observations may be reconciled by assuming that DNA binds MukB topologically (proposed model in Fig. S8).

Hydrolysis of ATP does only yield about 20 to 25 *k_B_T* energy. Actually, it was shown that 80% of the input energy of ATP was dissipated into heat and did not contribute to power the motor protein kinesin (*38*). The free energy released upon ATP hydrolysis is rapidly dissipated and it is the differential binding of ATP and its hydrolysis products ADP and Pi that leads to the slow conformational transitions of the motor protein (*39*). Numerous data has already shown that ATP hydrolysis is not involved in directional motion; rather, it drives SMC protein loading and unloading onto chromatin (*40, 41*). A relatively small force (entropic force) of *k_B_T/P* is required to align elastic units of dsDNA, and this entropic force is ~0. 1 pN, within the range of external forces that were applied in this work, and also published for single-molecule experiments that observed DNA compaction or looping induced by cohesion and condensin (*9*)(*11*) (*21*)(*42*). Moreover, in cells, DNA can move or slide by itself if considering the elastic energy that DNA contains, such as supercoiling (Fig. 3G) (*43*)(*44*)(*45*)(*46*).

Our DNA sliding model (Fig. 4F) does explain how MukBEF can efficiently organize DNA while having only low ATPase activity and the intrinsic ability to entrap DNA topologically and react as a ratchet. Nucleosomes containing ~150 base pairs of tightly wrapped DNA are the basic structural elements of chromatin compaction and regulation, while SMC proteins contribute to the large-scale chromosome organization and regulation. Nucleosomes do show sliding and loop formation. (*47*) (*48*) (*49*) (*50*)(*51*). Perhaps, SMC proteins and nucleosomes share a common DNA sliding, loop formation, and ratchet mechanism. Other SMC proteins can also be immobilized on the surface to test this DNA sliding and ratchet mechanism. How SMC complexes facilitate DNA chromosome organization and segregation touches fundamental biological and physical questions: What are the molecular mechanisms of motor proteins? What is the contribution of entropy in the living cells? Although we cannot rule out complete answers for these questions, our results suggest a possible molecular mechanism of SMC protein, which could rationalize all the single-molecule observations in this study.

## Materials and Methods

### Fabrication of flow cell

The glass slides and coverslips were super-cleaned with Acetone, 1M NaCl and MiliQ water solutions. The bottom coverslip was functionalized with an amino-group in the 2% 3-aminopropyltheithoxysilane (440140, Sigma) in acetone for 10 min. Then surfaces were PEGylated by applying a viscous mixture of PEG and biotin-PEG to one side of the coverslip at room temperature for 3 hours and washed intensively with MiliQ water.

Four 1.5 mm holes were drilled in the glass slide in the pattern show in Fig. 1B, and 1.5 mm diameter tubing was glued into each hole with epoxy resin glue. Afterwards, double-side tape was stack between coverslip and drilled glass slide to make a flow cell.

### Double-tethered *λ*DNA for single-molecule study

The double biotin-labelled *λ*DNA preparation was similar as ref.(*9*). Flow cells were first incubated with 0.3 mg/mL Neutravidin in T50 buffer (20mM Tris-HCl, pH=7.5, 50 mM NaCl) for 1 min and washed with 400 μl T50 buffer. 50 μl of 20 pM double biotin-labelled *λ*DNA (ThermoFisher Scientific; SD0011) was introduced into flow cells at 1 ~ 2 μl/min in the T50 buffer with 200 nM Sytox orange (ThermoFisher Scientific; S11368). Excess DNA was washed off with T50 buffer once optimum DNA density was achieved.

10-100 nM MukB-Cy5 in the crude image buffer without PCA/PCD and Trolox (20mM Tris-HCl, pH=7.5, 50 mM NaCl, 0.3mg/BSA, 2 mM DTT, 400 nM Sytox orange) was introduced at 1 μl/min, to observe DNA compaction. 10 nM MukF, 20 nM MukE and 1 mM ATP-Mg^2+^ in the imaging buffer (20 mM Tris-HCl, pH=7.5, 50 mM NaCl, 0.3 mg/BSA, 2 mM DTT, 2.5 mM PCA, 50 nM PCD, 1 mM Trolox and 400 nM Sytox orange) was introduced in the perpendicular direction.

For EQ mutation, 10 nM MukB^EQ^-Cy5, 10 nM MukF, 20nM MukE and 1 mM ATP-Mg^2+^ in the imaging buffer (20 mM Tris-HCl, pH=7.5, 50 mM NaCl, 0.3 mg/BSA, 2 mM DTT, 2.5 mM PCA, 50 nM PCD, 1 mM Trolox and 400 nM Sytox orange) was introduced into the flow cell, and then 10 nM MukF, 20 nM MukE and 1 mM ATP-Mg^2+^ in the imaging buffer (20 mM Tris-HCl, pH=7.5, 50 mM NaCl, 0.3 mg/BSA, 2 mM DTT, 2.5 mM PCA, 50 nM PCD, 1 mM Trolox and 400 nM Sytox orange) was introduced in the perpendicular direction.

For DA mutation, 100 nM MukB^DA^ was introduced in the flow cell.

### Immobilization of His_6_-tagged MukB for single-molecule study

Flow cell was first incubated with 0.3 mg/mL Neutravidin in T50 buffer (20 mM Tris-HCl, pH=7.5, 50 mM NaCl) for 1 min and washed with 400 μl T50 buffer. Afterwards, 2 g/ml Biotinylated Anti-His Antibody (Penta-His Biotin Conjugate; Qiagen; No. 34440) in T50 buffer was introduced into flow cells at 50 μl/min, then wash immediately with 300ul T50 buffer. 5 nM - 10nM MukB was incubated with 100 pM *λ*DNA or 100 pM 44kb plasmid in T50 buffer for 20 min at room temperature (22 °C), then introduced into the flow cell at 5-10 μl/min to immobilize His_6_-tagged MukB on the surface.

### Single-molecule Imaging

Single-molecule TIRF experiments were performed on a custom-built, objective-type TIRF microscope. A green laser (532-nm Cobolt Samba; Cobalt) and a red laser (635-nm CUBE; Coherent) were combined using a dichroic mirror and coupled into a fibre optic cable. The output of the fibre was focused into the back focal plane of the objective (magnification of 100×, oil immersion, N.A. = 1.4, f/26.5; UPlanSApo, Olympus) and displaced perpendicular to the optical axis such that laser light was incident at the slide/solution interface at greater than the critical angle, creating an evanescent excitation field. Illumination powers were set as low as possible to avoid the photo damage of DNA.

Fluorescence emission was collected by the objective and separated from the excitation light by a dichroic mirror (545 nm/650 nm; Semrock) and clean-up filters (545-nm long pass, Chroma; 633/25-nm notch filter, Semrock). The emission signal was focused on a rectangular slit to crop the image and then spectrally separated, using a dichroic mirror (630-nm long pass; Omega), into two emission channels that were focused side-by-side onto an electron multiplying charge coupled device (EMCCD) camera (iXon 897; Andor). The EMCCD was set to an EM gain of 300, corresponding to an approximate real gain of four counts per photon. Each pixel on the EMCCD corresponded to a 96 × 96-nm region in the imaging plane.

### Image analysis and data processing

Fluorescence images were analyzed by ImageJ software.

### Expression and purification of the MukBEF His_6_-tagged proteins

Wild-type MukB was 6×His-tagged at the C-terminus and was expressed from plasmid Pet21 in C3013I cells (NEB). For Immobilized His_6_-tagged MukB, a 10 amino acids flexible glycine-serine protein domain linker (or Halo Tag) was introduced between His tag and MukB. 2L cultures were grown in LB with appropriate antibiotics at 37 °C to OD600~0.6 and induced by adding IPTG at final concentration of 0.4 mM. After 4 hours at 30 °C, cells were harvested by centrifugation, suspended in 30ml lysis buffer (50 mM HEPES pH 7.5, 300 mM NaCl, 5% glycerol, 10 mM imidazole) supplemented with 1 tablet Complete mini-protease inhibitor cocktail (Roche) and sonicated. Cell debris was removed by centrifugation and target proteins were first purified by TALON Superflow resin. Then, the fractions from TALON were diluted to 100mM NaCl buffer and injected to HiTrapTM Heparin HP column (GE Healthcare) pre-equilibrated with Buffer A (50mM HEPES pH 7.5, 100mM NaCl, 5% glycerol, 1 mM EDTA, 1 mM DTT), then the column was washed at 1ml/min flow rate with a gradient 100-1000 mM NaCl. The eluted fractions were collected and dialyzed with dialysis buffer (25 mM HEPES pH 7.5, 25 mM KCl, 0.1 mM EDTA, 2 mM DTT, 1 mM PMSF, 5% Glycerol).

The 6×His-tagged MukE and MuF were also at the C-terminus. For MukE-6×His and MukF6×His purifications, fractions from TALON resin were diluted and injected into HiTrap DEAE FF column (GE healthcare) pre-equilibated in Buffer A, and then the column was washed at 1 ml/min flow rate with a gradient 100-1000 mM NaCl. The eluted fractions were collected and dialyzed with dialysis buffer. Protein concentration was estimated by UV absorption at 280 nm on Nanodrop and purity was confirmed by SDS-PAGE. Purified proteins were aliquoted, snap-frozen and stored at −80 °C until use.

### Expression, purification and fluorescent labeling of the MukB His_6_-tagged protein

To label MukB site-specifically, the unnatural amino acid p-azido-L-phenylalanine (AZP) was incorporated into the MukB (*52*). The plasmid pBAD-MukB-S718TAG-10GS-6×His was constructed as following: First, MukB gene was inserted into pBAD vector. MukB was 6×His-tagged at the C-terminus, and 10 aa flexible glycine-serine protein domain linker was introduced between His tag and MukB. Then an amber (TAG) codon was introduced at 718 position of hinge domain of MukB by site-Mutagenesis.

To express MukB incorporated with AZP, pBAD-MukB-S718TAG-10GS-6×His plasmid and a pEvol plasmid containing the engineered Amber suppressor tRNA/synthetase system carrying a chloramphenicol resistance marker, were transformed into a derived strain (FW01) from C321ΔA strain, with chromosomal MukB tagged with 3XFlag.

2L cultures were grown in LB with 100 μg/ml Carbanicillin, 25μg/ml chloramphenicol and 1% glucose at 30 °C to OD600~0.6 with 1mM p-azido-L-phenylalanine, and then induced by adding of L-arabinose at final concentration of 0.4% (W/V). After 4 hours at 30 °C, cells were harvested by centrifugation, suspended in 30ml lysis buffer (50mM HEPES pH 7.5, 300mM NaCl, 5% glycerol, 10mM imidazole) supplemented with 1 tablet Complete mini-protease inhibitor cocktail (Roche) and sonicated. Cell debris was removed by centrifugation and target proteins were first purified by TALON Superflow resin. Then, the fractions from TALON were diluted to 100mM NaCl buffer and injected to HiTrapTM Heparin HP column (GE Healthcare) pre-equilibrated with Buffer A (50 mM HEPES pH 7.5, 100mM NaCl, 5% glycerol, 1mM EDTA, 1 mM DTT), then the column was washed at 1ml/min flow rate with a gradient 100-1000mM NaCl. The eluted fractions. The chromosomal MukB was removed by adding ANTI-FLAG^®^ M2 Affinity gel (A2220, Sigma) into the eluted fractions. After incubation for 30 min at 4 °C, and the gel was discarded by centrifugation. The supernatant was collected and concentrated by ultrafiltration.

The purified protein was labelled via copper free click chemistry. The protein was incubated with ~20× molar excess of Dibenzylcyclooctyne(DBCO)-Sulfo-Cy5 (Jena Bioscience) at 4°C overnight in the dark. The Cy5 labelled MukB was purified by size-exclusion chromatography on a Superdex 200 10/300 GL column (GE Healthcare). Peak fractions were pooled and concentrated by ultrafiltration.

Labeling efficiency was estimated to be 65% based on the absorbance ratio of 280 and 650 nm, using the calculated molar extinction coefficient of MukB at 280 nm (335,670 M-1 cm-1), the molar extinction coefficient of the Cy5 dye at 650 nm (250,000 cm-1M-1.), and correct factor for the absorption at 280 nm by the dye (ε280 nm/ε650 nm = 0.05).

### Expression and purification of the MukE-Flag and MukF-Flag proteins

MukF Flag-tagged fragments were expressed from pET 21 plasmids in C3013I cells (NEB).The Flag-tagged MukE and MuF were at the C-terminus. pET21-MukE-C100S-Cys-Flag and pET21-MukF-C204S-Cys-Flag were constructed with the indigenous cysteines were removed and one cysteine was introduced at the C-terminus.

2L cultures were grown in LB with appropriate antibiotics at 37 °C to OD600~0.6 and induced by adding IPTG at final concentration of 1mM. After 4 hours at 30 °C, cells were harvested by centrifugation, suspended in lysis and binding buffer (50mM HEPES pH 7.5, 150mM NaCl, 10% glycerol, 1mM EDTA) supplemented with 1 tablet Complete mini-protease inhibitor cocktail (Roche) and sonicated. Cell debris was removed by centrifugation and target proteins were incubated with 2ml ANTI-FLAG^®^ M2 Affinity gel (A2220, Sigma) at 4°C for 4 hours. The Flag tagged proteins were eluted out with high salt buffer (50mM HEPES pH 7.5, 500mM NaCl, 10% glycerol, 1mM EDTA). The eluted fractions were collected and dialyzed with dialysis buffer. Purified protein was snap-frozen and stored at −80°C until use.

### ATP Hydrolysis Assays

ATP hydrolysis was analysed in steady state reactions using an ENZCheck Phosphate Assay Kit (Life Technologies). 150 μL samples containing standard reaction buffer supplemented with 2 mM of ATP were assayed in a 18 BMG Labtech PherAstar FS plate reader at room temperature. The results were computed using MARS data analysis software. Quantitation of phosphate release was determined using the extinction coefficient of 11,200 M-1cm-1 for the phosphate-dependent reaction at 360 nm at pH 7.0.

### Mutagenesis

Point mutations in all constructs were made by using Q5 Site-Directed Mutagenesis Kit (NEB). Primers were designed with NEBase Changer. 10 ng of the template was taken to the reaction. Plasmids were isolated and mutations confirmed by sequencing.

### Complementation assay

The assay relied on the ability of MukB expressed from pET21 in the absence of IPTG (leaky expression) to complement growth defect of ΔmukB (RRL149) cells at non-permissive temperature (37°C). Cells were transformed with pET21 carrying mukB or mukB variant and allowed to recover over night post transformation at permissive temperature (22°C) on LB plates containing Carbenicillin (100μg/ml) along with positive and negative controls. Then single colonies were streaked onto two fresh plates and incubated at the permissive or non-permissive temperature to assay their ability to complement the growth deficiency.

## AUTHOR CONTRIBUTIONS

M.Z and D.J.S conceived and directed the project. M.Z. undertook biochemical experiments, single molecule experiments and undertook quantitative imaging and analysis. M. Z. wrote the manuscript.

## ACKNOWLEDGEMENT

We thank Florence Wagner for plasmid preparation and EMSA assays. Gemma L. M. Fisher undertook the ATPase assay. Achillefs Kapanidis (Dept of Physics, University of Oxford) provided facilities for single molecule experiments and David Sherratt provided facilities for biochemical experiments in the Department of Biochemistry and directed some of the research. Members of Kapanidis group provided support for single molecule experiments. James Ross (University of Leeds) helped with crystal structure modelling. We thank David Sherratt and all members of the Sherratt lab for helpful discussions. We thank Achillefs Kapanidis and Jörg Enderlein (III. Institute of Physics of the Georg August University Göttingen) for insightful discussion and constructive criticism of the manuscript.

## Supplementary figures

**Fig S1.**
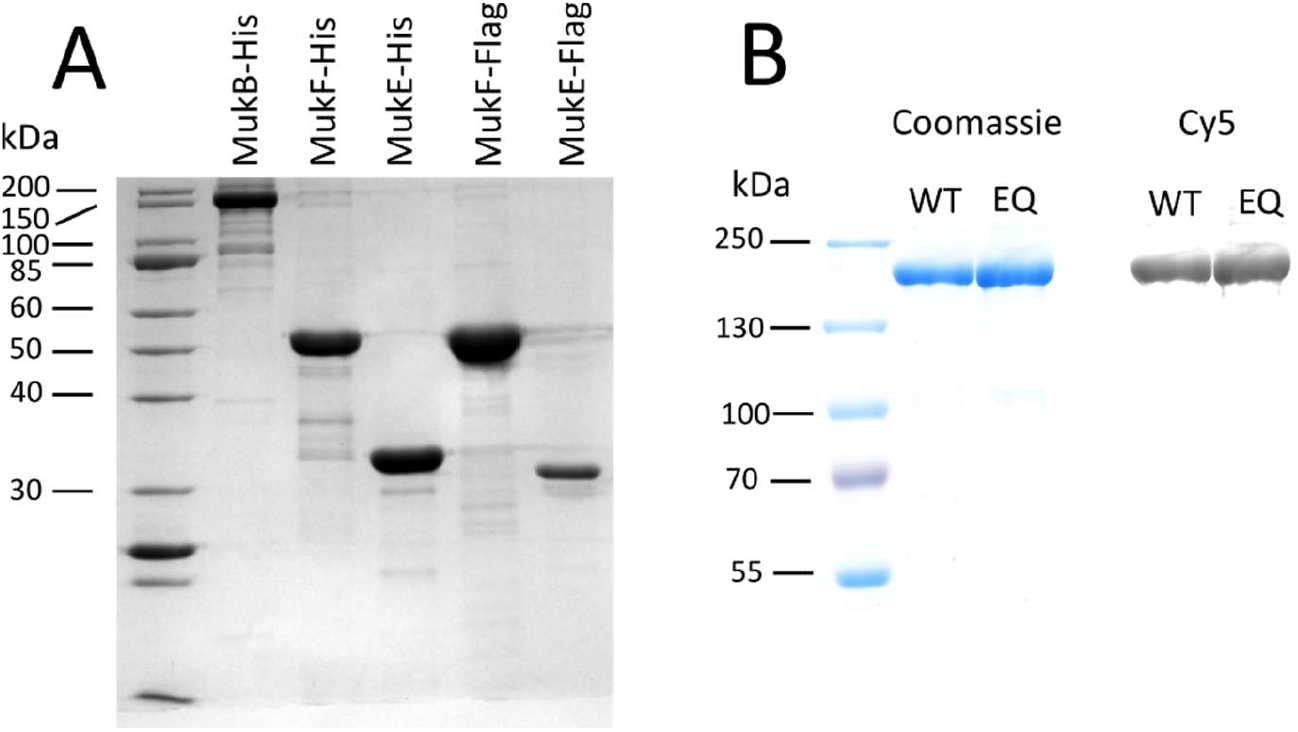
(A) Coomassie blue staining of purified recombinant E.coli MukBEF with His-tag, MukE-Flag and MukF-Flag. (B) The purified and labeled Wild-type MukB (WT) and ATP hydrolysis-defective mutant MukB^EQ^ (EQ) were analyzed by SDS-PAGE followed by Coomassie blue staining and in gel fluorescence detection of Cy5 dye. The labeling efficiency of homodimeric MukB is ~ 65%: a mixture of 35% unlabelled, 48.2% singly labelled, and 16.8% doubly labelled MukB is expected.

**Fig S2.**
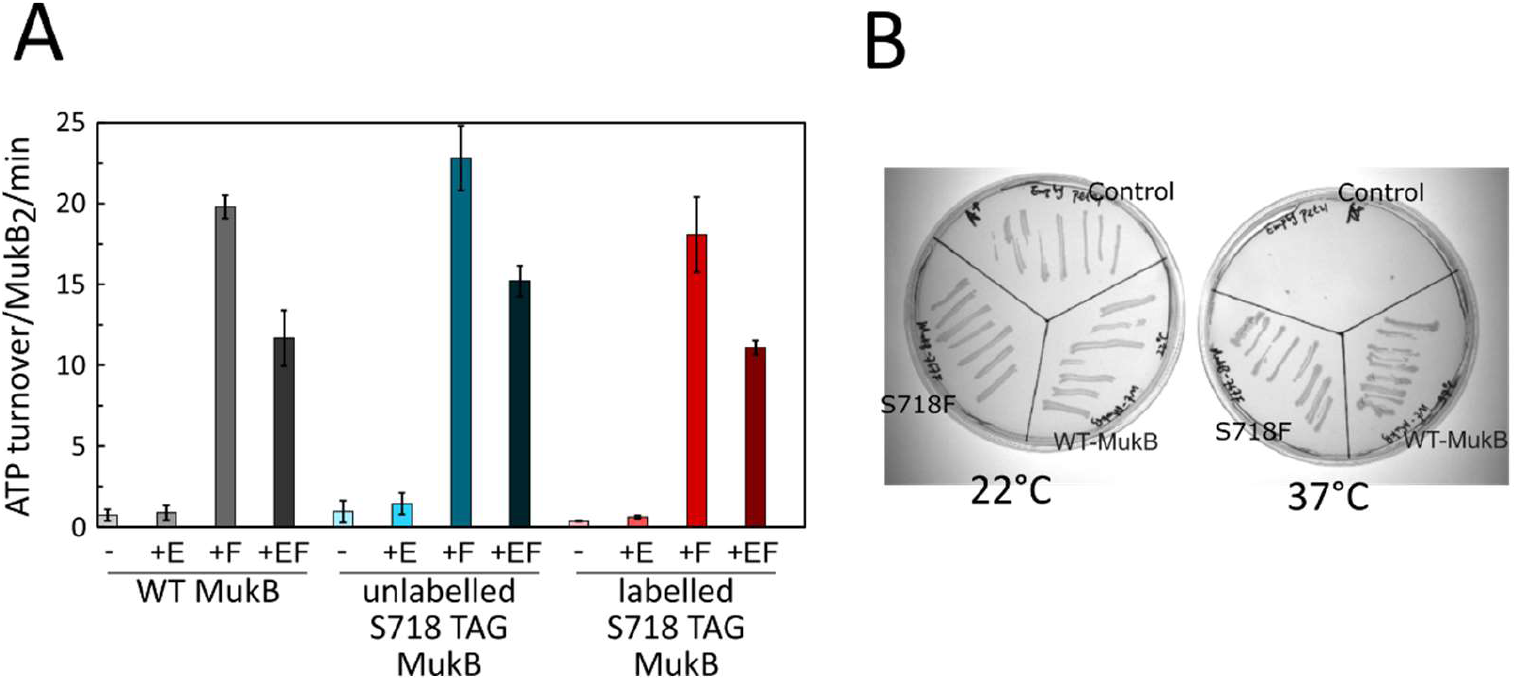
(A)The ATPase activities (Mean ± SEM) of MukB, S718TAG MukB, and Cy5(Cy3) labelled MukB, in the presence of the indicated components. (B) Functional analysis by complementation assay of mutated MukB_S718S. In vivo complementation in cells lacking chromosomal MukB gene ΔmukB (RRL149) by variants expressed from pET21 plasmid. Growth of material streaked of MukB_S718F was compared to growth of cells carrying WT-MukB construct at permissive, 22°C, and non-permissive, 37°C; Control: empty vector, a negative control. The MukB_S718F fully complements the temperature-sensitive growth defect of *ΔmukB* cells *in vivo*.

**Fig S3.**
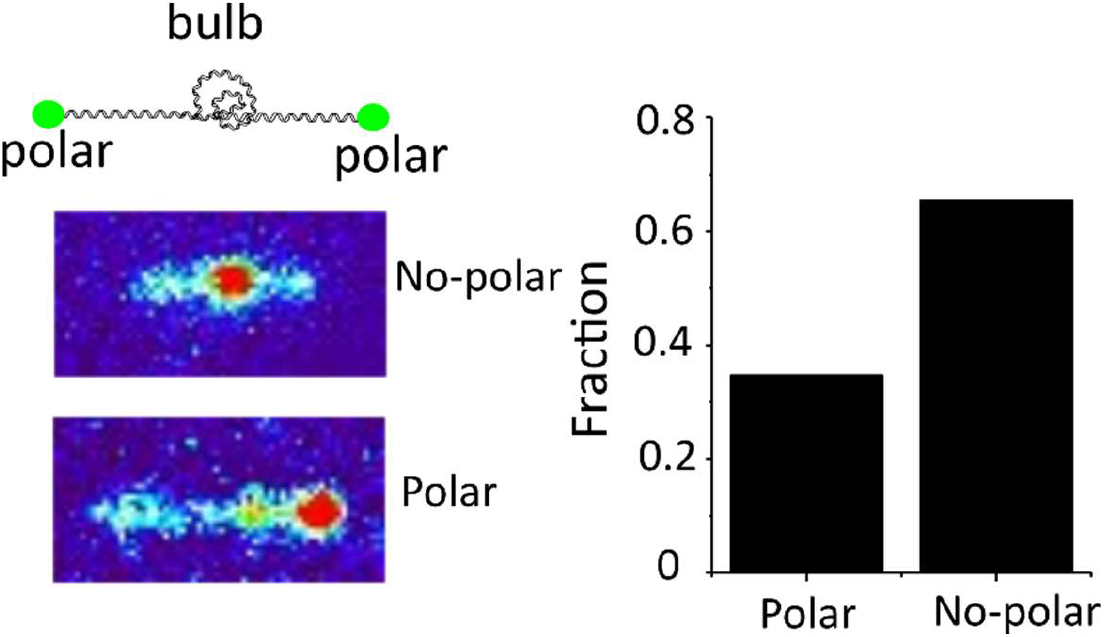
Fraction of DNA ‘bulb’ location. DNA ‘bulbs’ are located only by 34.6 % (83/240 DNA bulbs) at the polar regions of tethered *λ*DNA which indicates that ‘DNA bulb’ formation is not due to protein interaction between MukB and biotin/neutravidin on the surface.

**Fig. S4.**
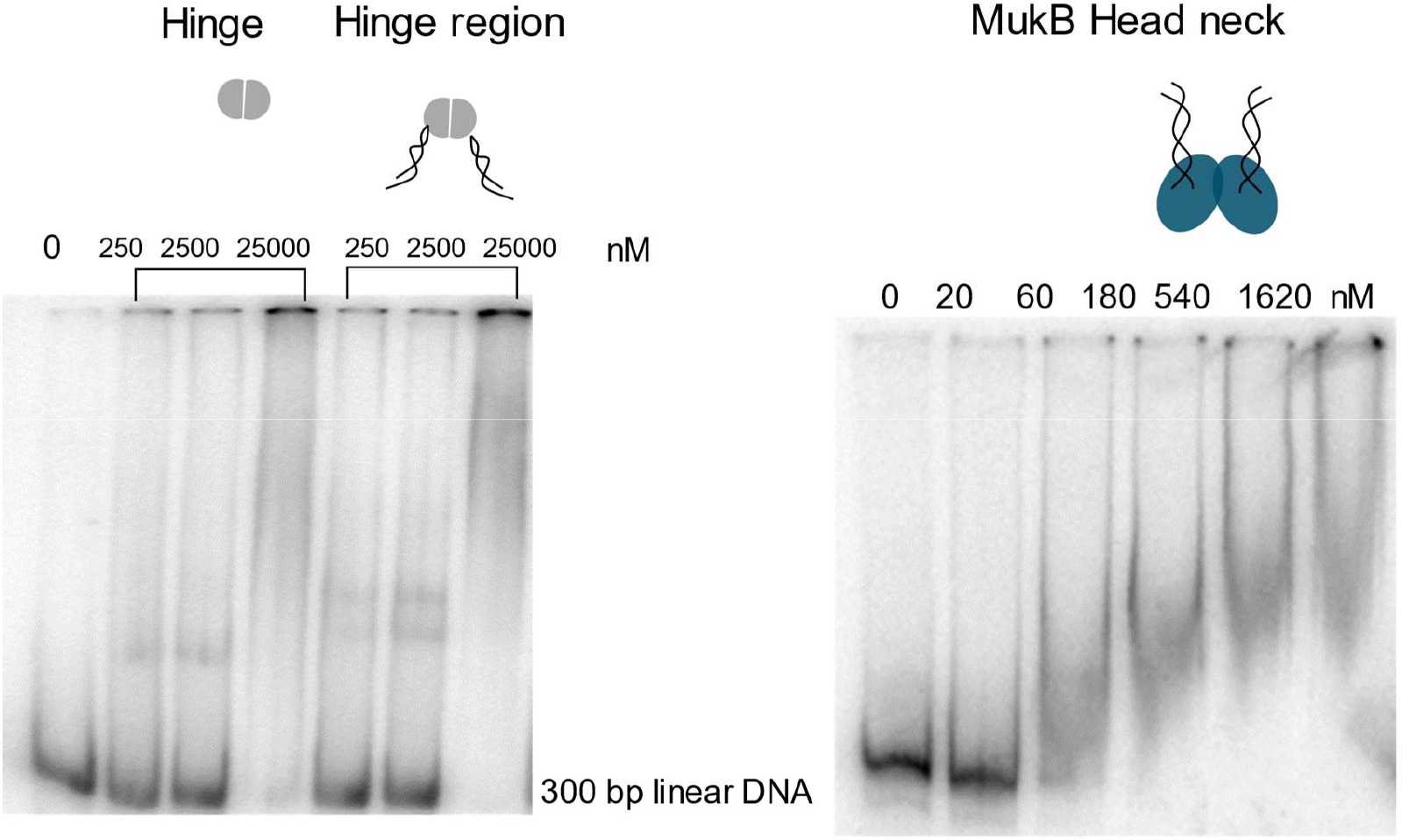
Gels from DNA electrophoretic mobility shift assay (EMSA) to test binding affinities of MukB fragments to DNA *in vitro*. The DNA fragment concentration was 300 nM. Hinge of MukB and Hinge range of MukB concentrations were: 250, 2500 and 25000 nM, respectively. MukB head neck concentration were: 20, 60, 180, 540, 1620 nM. Samples were incubated for 30 min at room temperature in T50 buffer, and applied to a 6 % polyacrylamide gel equilibrated with the same buffer. Electrophoresis was at 120 V for 1 h. Concentration “0” denotes free DNA. Very weak binding between DNA and MukB fragments can be observed.

**Fig S5.**
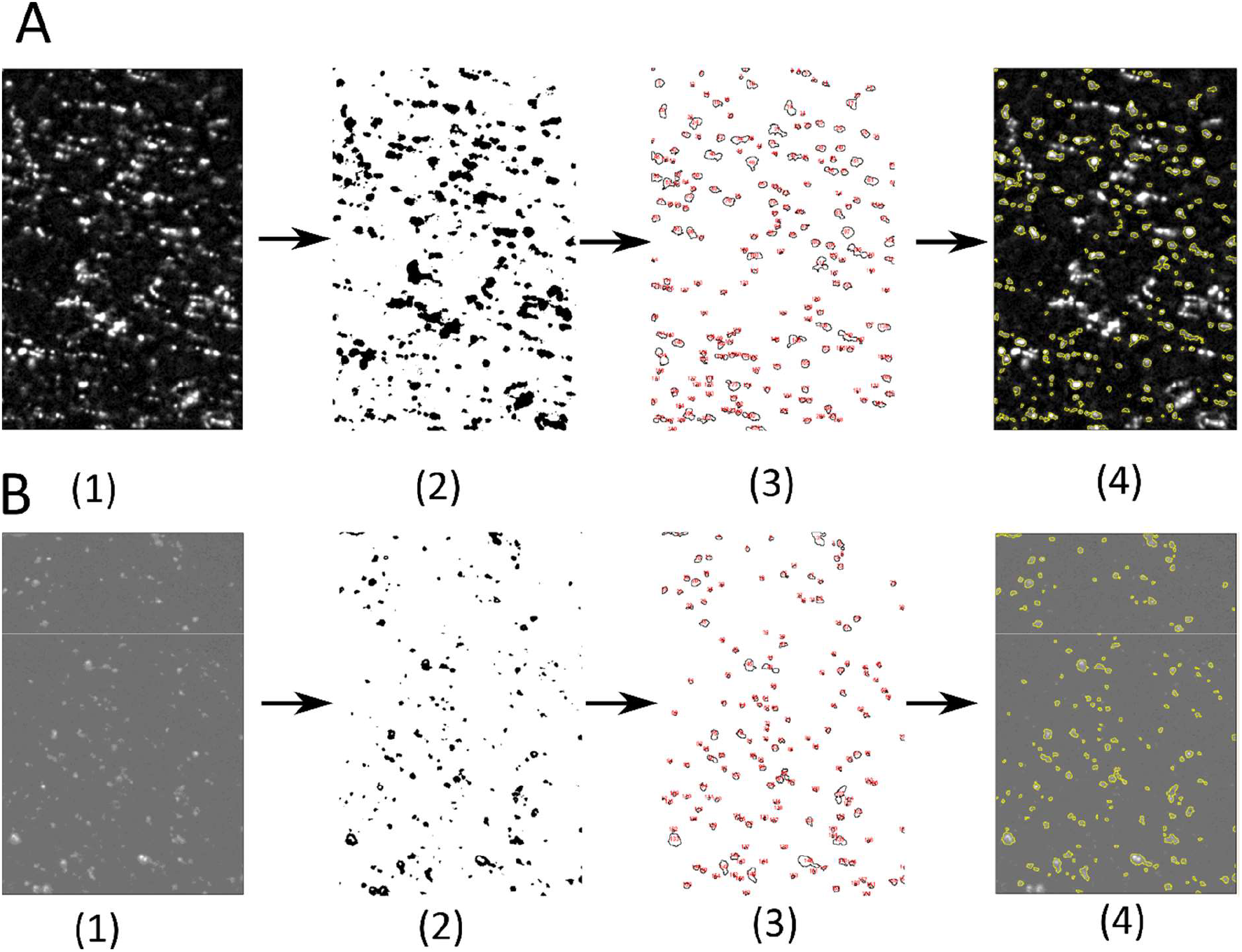
Example of imaging process to quantify intensity of MukB^EQ^ clusters with MukE and MukF (A) or MonoMukF (B) on the tethered DNA. (1) the image after subtracting background (2) A binary mask after applying thresholding to select the MukB^EQ^ molecules (3) identified ranges by applying ‘analyze particles’ on the mask, that are between 9-300 pixels and with a circularity of between 0.2 and 1. (4) overlapped structures with the original image. The intensity of selected ranges was calculated for number determination of MukB^EQ^.

**Fig. S6.**
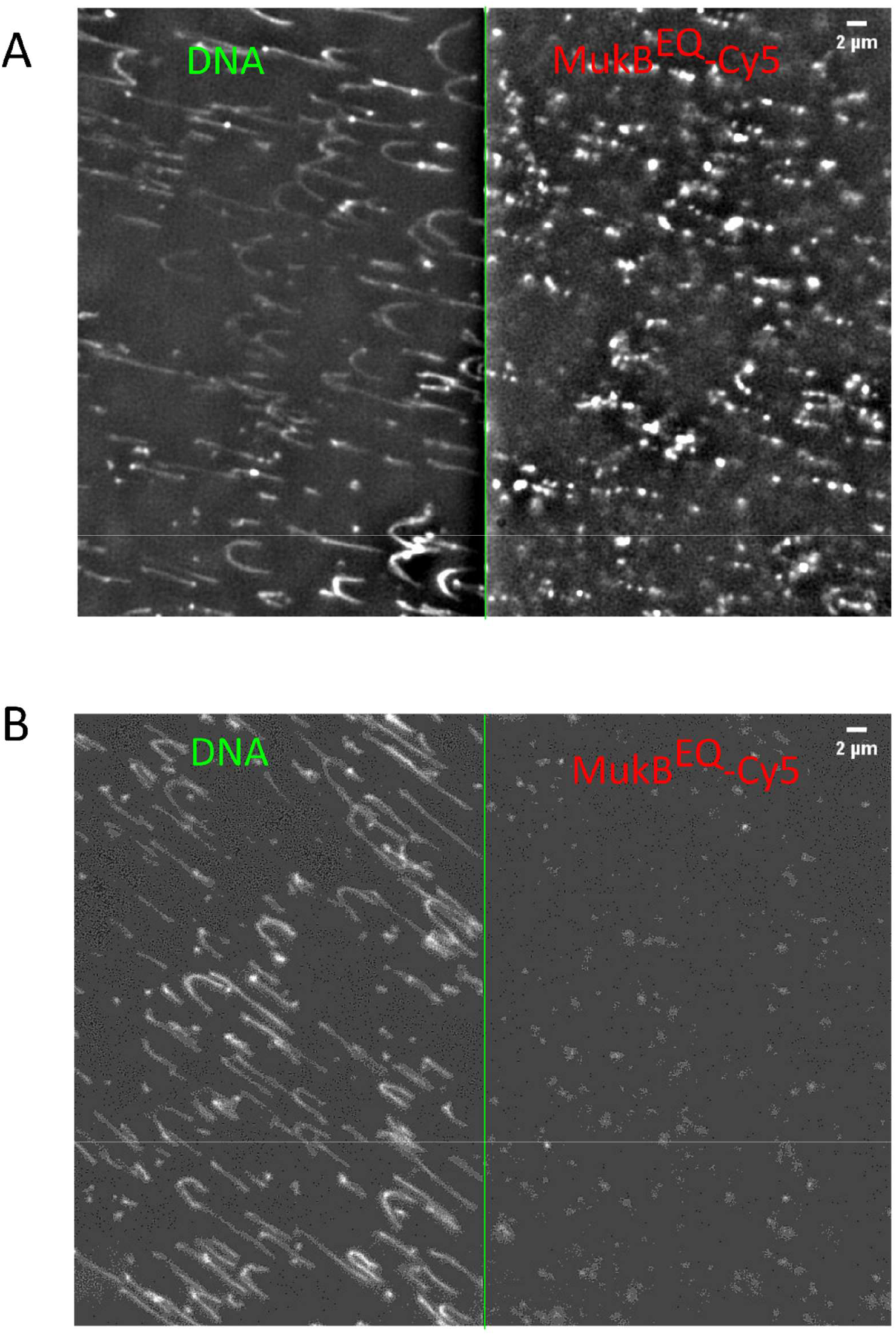
**(A).** Snap-shot of MukB^EQ^ EF clusters with ATP-Mg^2+^ in 150mM NaCl under 1 pN force. **(B).** Snap-shot of MukB^EQ^ with MukE, MonoMukF and ATP-Mg^2+^ in 150mM NaCl on the biotinylated anti-His_6_-antibody functionalized PEG surface. Left channel is Sytox Orange stained DNA, and the right channel is Cy5 labelled MukB^EQ^.

**Fig. S7.**
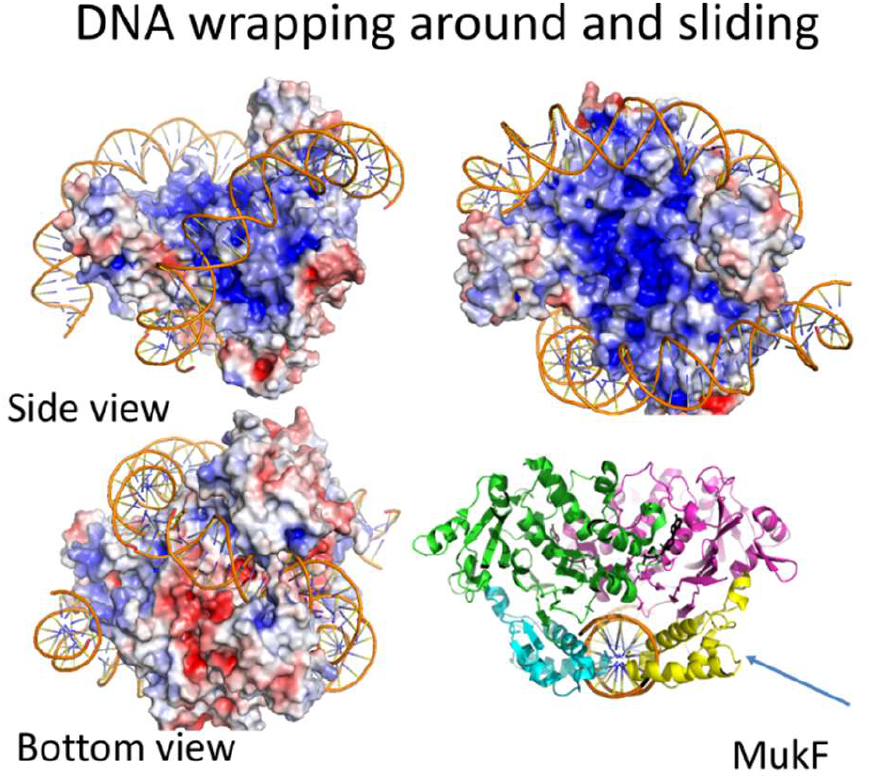
DNA wrapping around the head domain of MukB and sliding. Proposed model based on published crystal structure (Woo et al., 2009): DNA is topologically entrapped by a MukB head in the presence of the MukF C terminus. C terminus of MukF inserts into the groove of DNA (also see video 14). Structural studies of wide-type SMC complex would help verify the DNA binding sites. This DNA sliding model can also explain the different behaviours of other SMC-similar proteins. For example, Mre11-nuclease & Rad50-ATPase complex show different properties when interacting with linear or with circular DNA (not supercoiled), while linear DNA is more probable to slide into the Rad50 ring to allow digestion by Mre11-nuclease (*53*). Also, cohesion complexes act differently in the presence of linear and circular DNA (not supercoiled) attached to beads under high salt concentration, where a cohesion complex can slide off the free ends of linear DNA, but gets trapped on circular DNA (*54*).

**Fig. S8.**
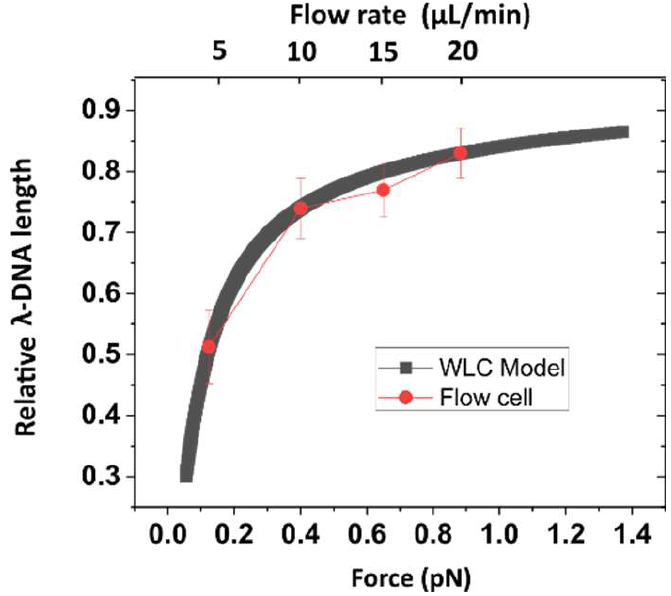
Calibration of applied force and flowrate. The flowrate is between 1ul/min to 100ul/min. The applied force is calibrated based on the previous studies (*55*).The relationship between applied flow rate and relative *λ*DNA length, fitted to the Worm-Like Chain (WLC) model (solid line) with the fitted persistence length of 43 nm and a contour length of 16.3 μm.

